# Protein Assembly Modulation: A New Approach to ALS Therapeutics

**DOI:** 10.1101/2023.07.24.550252

**Authors:** Shao feng Yu, Kumar Paulvannan, Dennis Solas, Anuradha F. Lingappa, Ana Raquel Moreira, Shriya Sahu, Maya Michon, Danielle Goldsmith, Nicholas DeYarman, Suguna Mallesh, M. Dharma Prasad, Claudia Maios, Kai Ruan, Giulio S. Tomassy, Elizabeth Jensen, Emma McGuirk, Verian Bader, Andreas Mueller-Schiffmann, Jonathan C. Reed, Jaisri R. Lingappa, Vinod Asundi, Shi Hong, Steve Jacobsen, Lyle Ostrow, Tom Lloyd, Alex Parker, Kim A. Staats, Justin Ichida, James Dodge, Debendranath Dey, Carsten Korth, Suganya Selvarajah, Vishwanath R. Lingappa, Jeffrey Rosenfeld

## Abstract

Amyotrophic Lateral Sclerosis (ALS) is a neurodegenerative disease with a complex, multifactorial pathophysiology, most commonly manifest as loss of motor neurons. We introduce a new mechanism of ALS pathogenesis via a novel drug-like small molecule series that targets protein disulfide isomerase (PDI) within a previously unappreciated transient and energy-dependent multi-protein complex. This novel drug was found to have activity in cellular models for both familial and sporadic ALS, as well as in transgenic worms, flies, and mice bearing a diversity of human genes with ALS-associated mutations. These compounds were initially identified as modulators of human immunodeficiency virus (HIV) capsid assembly in cell-free protein synthesis and assembly (CFPSA) systems, with demonstrated antiviral activity in cell culture. Their advancement as ALS-therapeutics, and the subsequent separation of activity against HIV and ALS in chemical subseries through structure-activity-relationship optimization, may provide insights into the molecular mechanisms governing pathophysiology of disordered homeostasis relevant to ALS.

## Background

Aberrant protein aggregation is a common pathophysiologic mechanism implicated in a variety of neurodegenerative disorders (1,2). In the case of ALS, a serious neurodegenerative condition primarily involving motor neurons, it is generally accepted that a cellular manifestation of disease is the mislocalization and aggregation of the protein transactive DNA-binding protein of 43 kDa (TDP-43) (3–8). In healthy individuals, TDP-43 is localized to the cell nucleus (9,10). However, at autopsy, almost all cases of ALS have shown TDP-43 mislocalized to the cytoplasm to varying degrees (5,6,9,10). In these cases, the mis-localized TDP-43 is found in aggregates, typically co-localized with stress-granule proteins (7,11,12). For poorly understood reasons, the end result is selective death of motor neurons.

ALS is a challenging disease to study, diagnose, and treat because it is heterogenous in phenotype and progression, both in cells and clinically, in patients (13–15). The overwhelming majority of ALS cases are sporadic, meaning that the ALS patient does not have a clearly identifiable genetic cause or a family history of the disease (16). Among sporadic ALS cases, age of onset, manifestation, and disease progression are variable. While in some cases patient’s symptoms worsen quickly, in another subset they progress slowly (13,15). A small subset of ALS is familial, for which specific gene mutations have been identified (16–18). The proteins encoded by these genes, and those with which they interact, comprise the ALS interactome (5,19). The specifics of how exactly these gene products work together normally, or malfunction to cause ALS, have not been established (20). Likewise, specific toxins have been implicated in the increased incidence of ALS-like syndromes, but their significance for sporadic ALS remains unknown (21).

The identification of specific genes in familial ALS has made possible the construction of transgenic animal models that show key phenotypic manifestations of ALS. *Caenorhabditis elegans* is a simple model with a total of 302 neurons (22). The *C. elegans* models for ALS have human TDP-43 or FUS gene mutations or C9orf72 repeat expansions which show neuronal degeneration (23). Wildtype *C. elegans* are able to swim in liquid media, but when human ALS-causing mutant transgenic *C. elegans* are placed in liquid medium they display swimming-induced paralysis likely in response to stress (23). *Drosophila melanogaster* is a more complex animal model with nearly three orders of magnitude more neurons (24). *D. melanogaster* with human C9orf72 repeat expansion transgenes show retinal neurodegeneration and developmental lethality (25). Mouse models for ALS, where TDP-43 mutations, C9orf72 repeat expansions, or SOD mutations are introduced as transgenes, show paralysis and neurodegeneration (26).

Viral capsid formation is perhaps the most robust protein assembly pathway known(27). Long viewed as occurring through spontaneous self-assembly(28), a body of literature suggests kinetic control over that thermodynamic endpoint of capsid formation through host-mediated catalysis(29–31). We hypothesized that protein aggregation diseases, including ALS, might be related to disordered protein assembly(32). If viral capsid formation is host-catalyzed, then perhaps that is the case more generally for protein assembly. It is only a small further extension of the hypothesis to suggest that protein aggregation diseases reflect dysregulation of a normal catalyzed assembly event. In the case of viral infection the process of repurposing host machinery has been fine-tuned through evolution. In the case of our hypothesis on protein aggregation diseases that process would be more stochastic but once occurring would be just as inexorable. A corollary is that drugs restoring homeostasis by targeting host machinery involved in catalyzed capsid assembly may also be therapeutic for diseases of protein aggregation, such as ALS, if they represent dysfunction of the same or related machinery.

We utilized cell-free protein synthesis and assembly (CFPSA) systems to define a catalyzed assembly pathway for viral capsid formation(33,34). That system was then adapted into a moderate throughput drug screen to identify small molecules that block formation of viral capsids (29,30,35,36). We screened a 150,000 compound library for small molecules with protein assembly modulating properties with respect to capsid formation for each of the viral families causing significant human disease, identifying a small numbers of hit compounds for each(27). These “protein assembly modulator” compounds were validated against infectious viruses (29,30,35–37). Subsequently, they were tested in cellular and animal models for nonviral disease, with success (35,37). Data from the application of assembly modulation to oncology indicates that defects in protein assembly are points of overlap in the molecular-level departures from homeostasis that drive progression of both viral and neoplastic disease (35). Treatment with assembly modulators appears to change the composition of particular multi-protein complexes which are comprised of a number of proteins implicated in pathogenesis of the disease, and others implicated in restoration of homeostasis, including through autophagy (35,36). One class of antiviral assembly modulators was shown to target the allosteric regulator 14-3-3 while a second class of assembly modulators with anti-cancer activity was shown to target the allosteric regulator KAP1/TRIM28 (35). We hypothesized that allosteric sites on these catalytic multi-protein complexes could be modulated by our compounds in ways that reverse the disease-associated changes and restore homeostasis, and thereby could have relevance in ALS models associated with aberrant protein aggregation.

Applications of these antiviral compounds in the realm of neuroscience in general and ALS in particular, was prompted by two additional considerations. First, that there is a long and enigmatic history of association of particular viruses with specific neurodegenerative diseases. Thus influenza has long been associated with Parkinson’s Disease, herpesvirus infections with Alzheimer’s Disease, and endogenous retroviral activation is observed in ALS (32,38–42). Indeed, the emergence of cognition through natural selection may have been due in substantial measure to retroviral-mediated genetic novelty (43). Second, endogenous retroviral activation has been associated with ALS(42). Together with the forementioned hypothesis that the protein aggregation observed in neurodegenerative disorders may be a variation on the theme of protein assembly, potentially reversible with assembly modulation (32,37), these considerations provided biologically plausible rationales for the line of investigation pursued here. Specifically, in view of these considerations, we used protein assembly modulator compounds active against capsid assembly of retroviruses including HIV as the starting point for our ALS counter screen (30,44). The results, reported here, provide a new framework for understanding the underlying pathophysiology of ALS. We show that a chemical series originally identified and validated against HIV, apparently also corrects a molecular-level defect responsible for TDP-43 mislocalization thereby restoring homeostasis in multiple models of ALS. Importantly, this chemical series can be progressed to a subseries lacking anti-HIV activity with enhanced therapeutic potency for treatment of ALS. Thus, the anti-viral target and the anti-ALS target, while related, appear distinct.

## Results

### Activity of HIV-assembly modulating compounds in cellular models of ALS

A phenotypic screen was established for identifying drug-like small molecule compounds which inhibited HIV capsid assembly in a CFPSA system (30). Hit compounds were termed “protein assembly modulators” due to their observed properties in of blocking capsid formation in CFPSA and cellular systems (30,35,36). The antiviral activity of some hit assembly modulators was validated against infectious HIV in cell culture (30,35). Some validated anti-HIV molecules were members of a tetrahydroisoquinolone (THIQ) chemical series (see **Supplemental Figure 1A** for structure and synthetic scheme of PAV-073, an advanced ALS-selective molecule within this series) (30).

Fibroblast cells from patients with familial ALS mutations and from sporadic frontotemporal dementia (FTD) patients, as well as healthy controls, were immunostained for TAR DNA-binding protein 43 (TDP-43) which is implicated in both retroviral infection and ALS pathology (36). With high content imaging of these immunostained patient-derived fibroblasts (PDFs), the mislocalization of TDP-43 from the nucleus to the cytoplasm is visible in sporadic, TDP-43 mutant, and VCP mutant fibroblasts, but not the healthy fibroblasts (see **Figures 1A** and **1B**). The cytosolic TDP-43 could be observed relocalized to the nucleus upon treatment with active THIQ compounds (see **Figures 1A** and **1B**).

**Figure 1.**
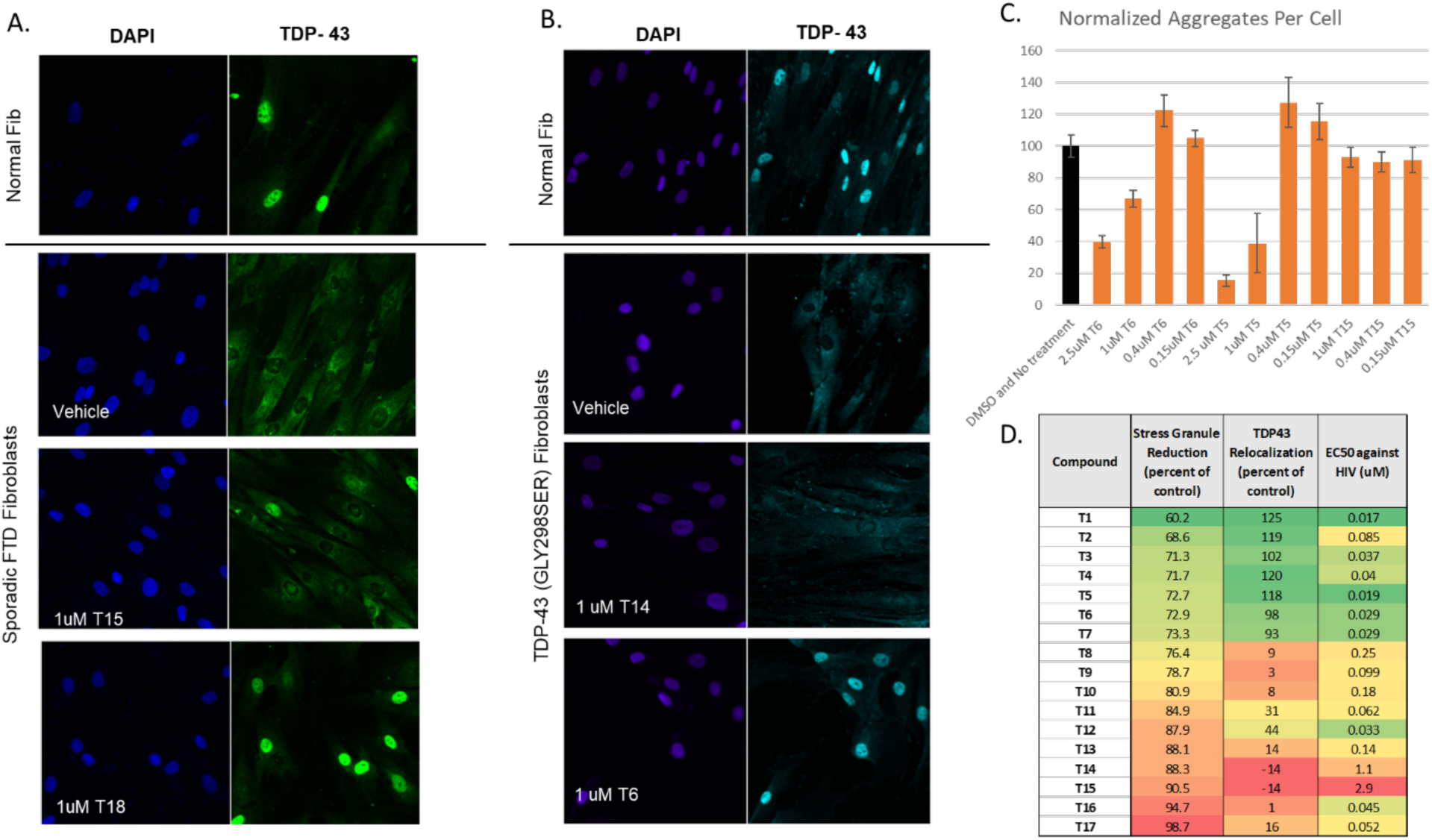
Activity of THIQ assembly modulators in cellular disease models. Figures 1A and 1B show TDP-43 mislocalization and relocalization upon treatment with active compounds for PDFs derived from ALS patients and healthy controls. In the nucleocytoplasmic relocalization assay, cells were seeded and treated with vehicle or compound for 4 days then washed, fixed, permeabilized and immunostained for TDP-43 and DAPI. Figure 1C shows quantitation of stress granule reduction in PDFs following treatment with compound. In the stress granule reduction assay, PDFs were treated with compound or vehicle for 24 hours then treated with 500uM sodium arsenite for one hour. Arsenite was washed off and cells were fixed, permeabilized, and immunostained for TDP-43, Hur, and DAPI. Cell profiler imaging was used to calculate the number of TDP-43 positive HuR aggregates per cell under each condition and those values were graphed. Figure 1D shows side-by-side comparison of values from the quantitation of the stress granule reduction assay, the nucleocytoplasmic relocalization assay, and activity against infectious HIV for chemical analogs within the THIQ series. In the infectious virus assay, MT-2 cells were infected with NL4-3 Rluc HIV and treated with compound or vehicle for four days. Anti-viral activity is shown as the calculated EC50.

**Figure 2.**
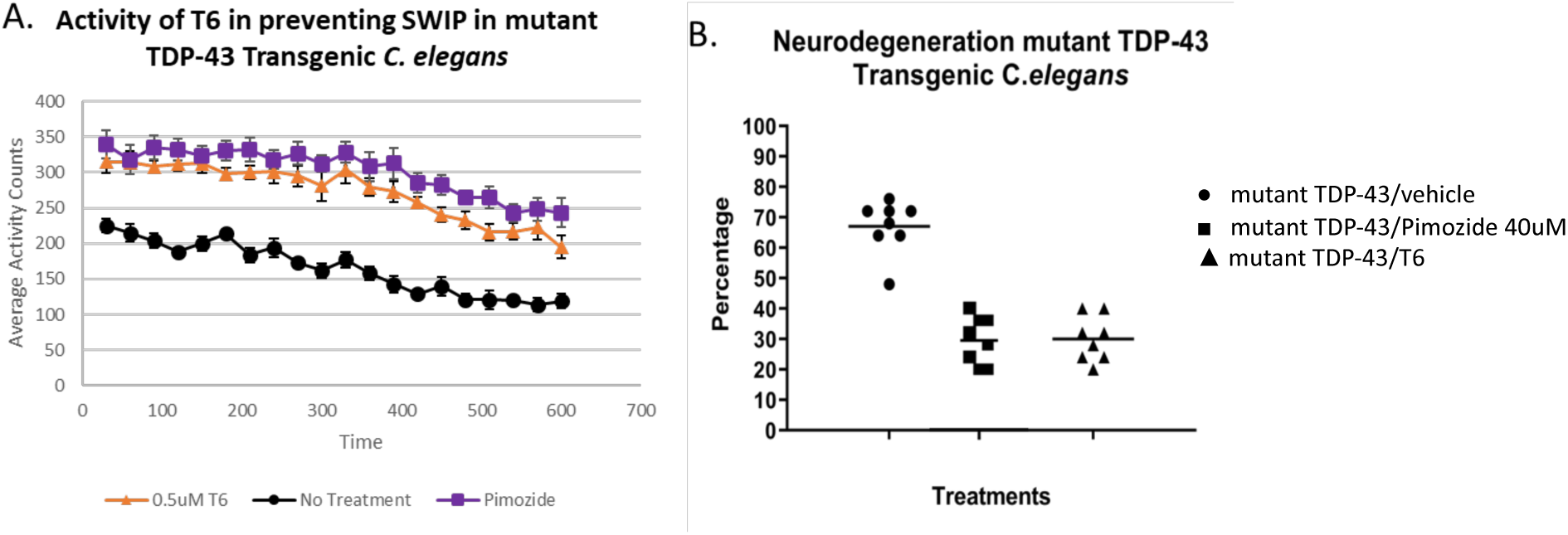
Activity in the *C. elegans* model for ALS. Figure 2A shows rescue of *C. elegans* transgenic for the TDP-43 A315T mutation in the SWIP assay. Worms were grown in the presence of 0.5uM THIQ compound T6, 40uM pimozide, or nothing and movement was recorded by video. The experiment was performed in 8 replicates of 25 worms each and the average number of body bends per second is shown for each condition. Figure 2B shows rescue of *C. elegans* transgenic for the TDP-43 A315T mutation in the long term neurodegeneration assay. Worms were grown in the presence of 0.5uM THIQ compound T6, 40uM pimozide, or vehicle for 9 days and analyzed for motor neuron splits. Percentage of neurons showing splits is shown for each condition.

**Figure 3.**
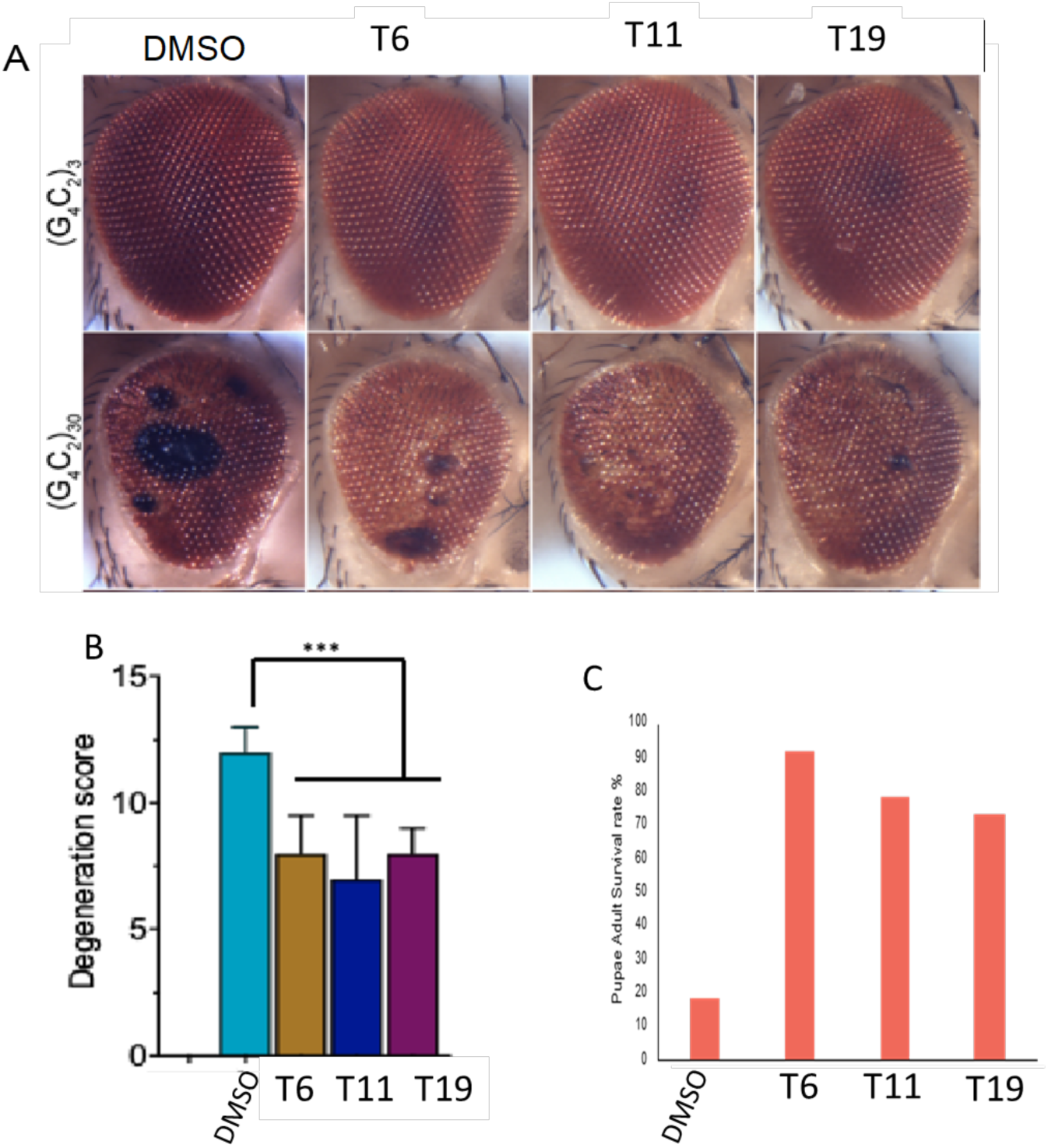
Activity in the *D. melanogaster* model for ALS. THIQ compounds T6, T11, and T12 were tested alongside vehicle in wildtype and transgenic *D. melanogaster* overexpressing C9orf72 30 G4C2. Figure 3A shows images of degeneration (black spots) in drosophila eye for wildtype (top row) and transgenic (bottom row) animals. Figure 3B shows the corresponding quantitation for degeneration observed in drosophila eye. Figure 3C shows percent adult survival in vehicle versus compound treated conditions.

**Figure 4.**
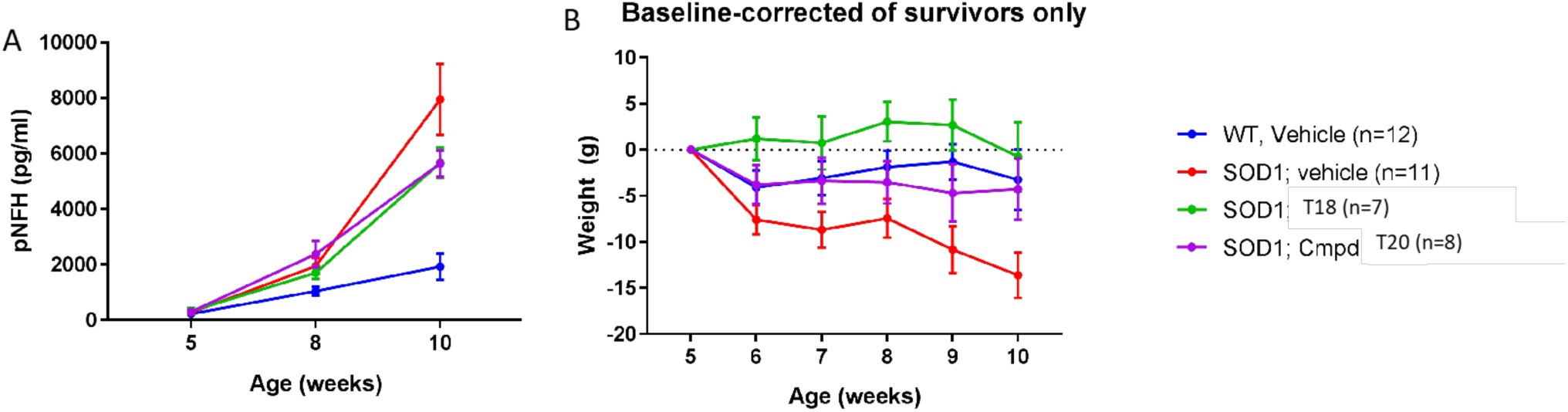
Activity in the SOD1 mouse model for ALS. THIQ compounds T18 and T20 were tested alongside vehicle in transgenic mice expressing the SOD G93A mutation. Figure 4A pNFH and Figure 4B shows weight over the course of the study with significant improvements in the compound-treated animals. Error bars represent standard deviation.

As a second cellular model for ALS, when PDFs were treated with 500 uM sodium arsenite for one hour and immunostained, cytoplasmic TDP-43 was found co-localized in stress granules along with HuR (see **Supplemental Figure 2**). Treatment with compounds active in the nucleocytoplasmic assay eliminated the stress granules in a dose-dependent manner (see **Figure 1C**). Following the SAR, most active compounds from the nucleocytoplasmic localization assay also displayed activity in the stress-induced stress granule assay (see **Figure 1D**).

The structure activity relationship (SAR) of early compounds from the THIQ series in the nucleocytoplasmic relocalization assay in sporadic FTD fibroblasts generally correlated to their activity against infectious HIV in MT-2 cells (see **Figure 1D**). However, the ALS and HIV activities were separable with further medicinal chemistry advancement, where some analogs (ex. compound T16) showed no TDP-43 relocalization but retained activity against HIV in the nanomolar range, while others (ex. compound T8) showed strong TDP-43 relocalization but substantially weaker antiviral activity compared to other potent compounds (see **Figure 1D**). Similarly, progression of the THIQ lead series resulted in a moderation of toxicity (see **Supplemental Figure 3)**. Precisely such a correlation of lowered toxicity to target selectivity for the virally modified multi-protein complex has been observed with lead series advancement for a structurally unrelated respiratory viral capsid assembly modulator(36).

### Activity of assembly modulating compounds against animal models of ALS

After achieving efficacy in multiple cellular models for familial and sporadic ALS, we turned to animal models. Lead compounds of the THIQ series were assessed in *C. elegans* with transgenic human ALS TDP-43 A315T mutation. In the swimming induced paralysis (SWIP) assay, transgenic worms were placed in liquid media containing vehicle or compound at a particular concentration. Worms in the liquid media were scored as “paralyzed” if their body cannot make a bending “S” movement. Efficacy was measured as average body bends per second in vehicle-treated versus compound-treated populations. In the neurodegeneration assay, transgenic worms were grown for 9 days and analyzed for motor neuron splits in the presence of vehicle or compound(45)(23). THIQ compounds demonstrated significant reduction of SWIP and long-term neurodegeneration.

Active compounds from the series were then assessed in *D. melanogaster* transgenic for the C9orf72 30 G4C2 repeat expansion. Overexpression of 30 G4C2-repeats in fly motor neurons using OK371-GAL4 causes lethality due to paralysis, preventing the exclusion of the adult from the pupal case. Treatment significantly reduced lethality and degeneration caused by 30 G4C2 repeats than when compared with DMSO vehicle alone.

We then wanted to assess activity in a mouse model. Vehicle or compound was administered to mice transgenic for the SODG93A mutation every day for 5 weeks (beginning when the mice were 5 weeks old) by intraperitoneal (IP) dosing. Vehicle was also administered by IP to wildtype mice as a control. Both test compounds showed significant, positive results based on biochemical data (levels of phosphorylated neurofilament heavy chain subunit measured in plasma) and clinical criteria (prevention of weight loss).

### Target identification in ALS and FTD patient fibroblasts and mouse brain samples

Previous studies on protein assembly modulator small molecule mechanism of action has shown that they target dynamic multi-protein complexes, a feature which appears shared by structurally-unrelated assembly modulator chemotypes efficacious in other therapeutic areas (35,36). The formation and action of these multi-protein complexes appears to be dependent on metabolic energy (nucleotide triphosphate hydrolysis). Protocols for energy-dependent drug resin affinity chromatography (eDRAC) provided a method to characterize the targets of assembly modulating compounds (35,36). In those experiments, extract from a disease-relevant cell line or tissue sample would be incubated with a modified analog of a compound attached to an Affi-gel resin and serve as an affinity ligand for target identification (46). The eDRAC experiments made possible tandem mass spectrometry (MS-MS) determination of protein composition of the isolated target multi-protein complexes under various conditions including healthy versus disease cells/tissues, with and without metabolic energy supplementation, and under vehicle versus compound treatment conditions (35,36). We sought to apply the same techniques to the ALS-active assembly modulators in order to better understand their targets and mechanism.

Cellular extract was prepared from the brain tissue of a wildtype mouse and a transgenic mouse expressing the SOD1 G93A mutation. Extract was supplemented with an “energy cocktail” of ribonucleotide triphosphates (to a final concentration of 1mM rATP, 1mM rGTP, 1mM rCTP, 1mM UTP), creatine phosphate, and 5 ug/mL creatine kinase. Extract was incubated on an Affi-gel resin coupled to a compound from the THIQ series which exhibited potent activity in both SGA, NCA, or a control resin which consisted of an affi-gel matrix couple to itself, for an hour at 22°C (30). The resins were washed with 100 bed volumes of buffer and eluted with 100 uM compound containing the energy cocktail first for two hours at 22° C, and then a second eluate collected overnight, followed by stripping the column with SDS. The overnight eluate was analyzed by MS-MS. 166 proteins were identified by spectral count in the wildtype eDRAC eluate (see **Figure 5A**). 208 proteins were identified by spectral count in the SOD1 eDRAC eluate (see **Figure 5A**).

**Figure 5.**
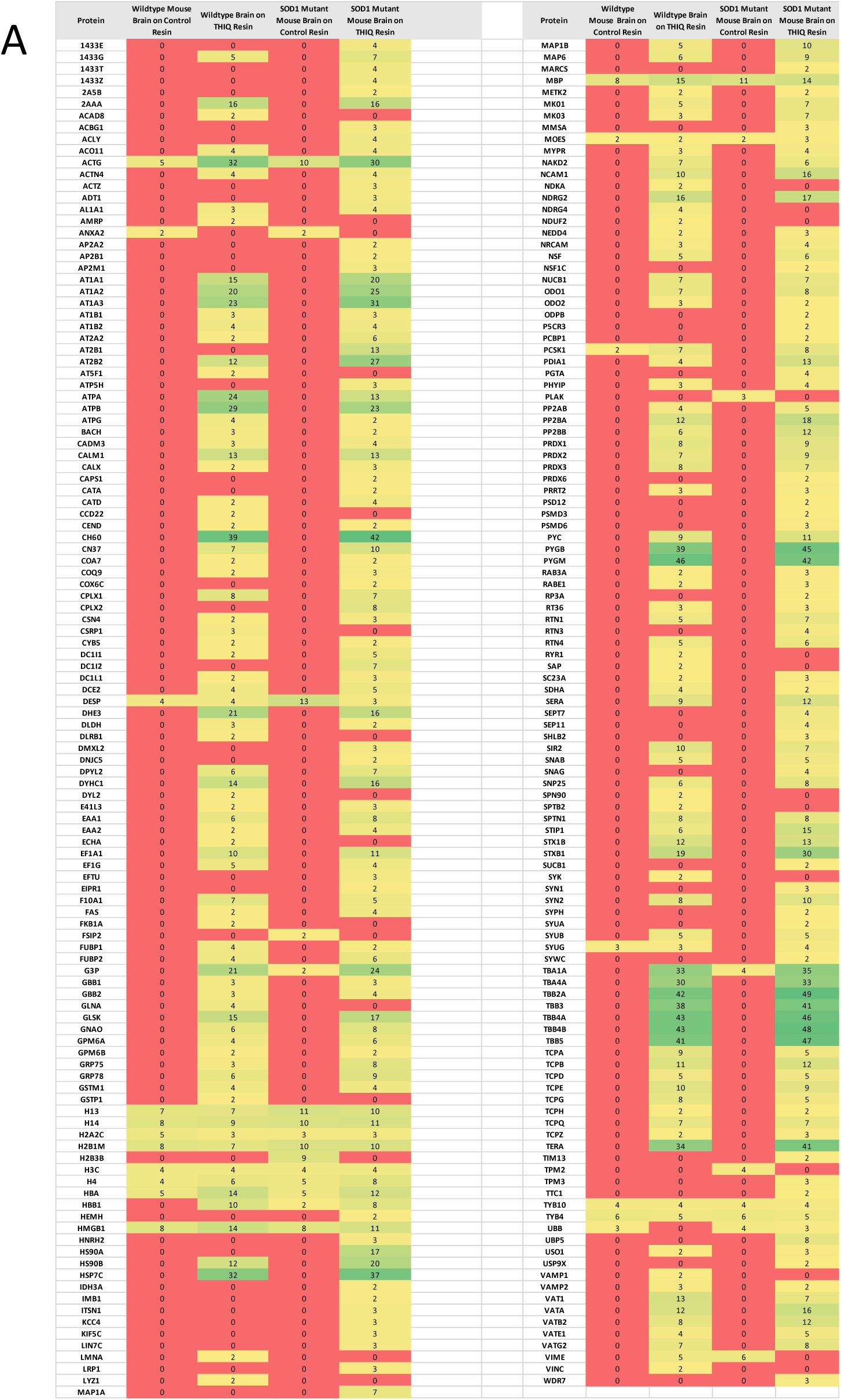
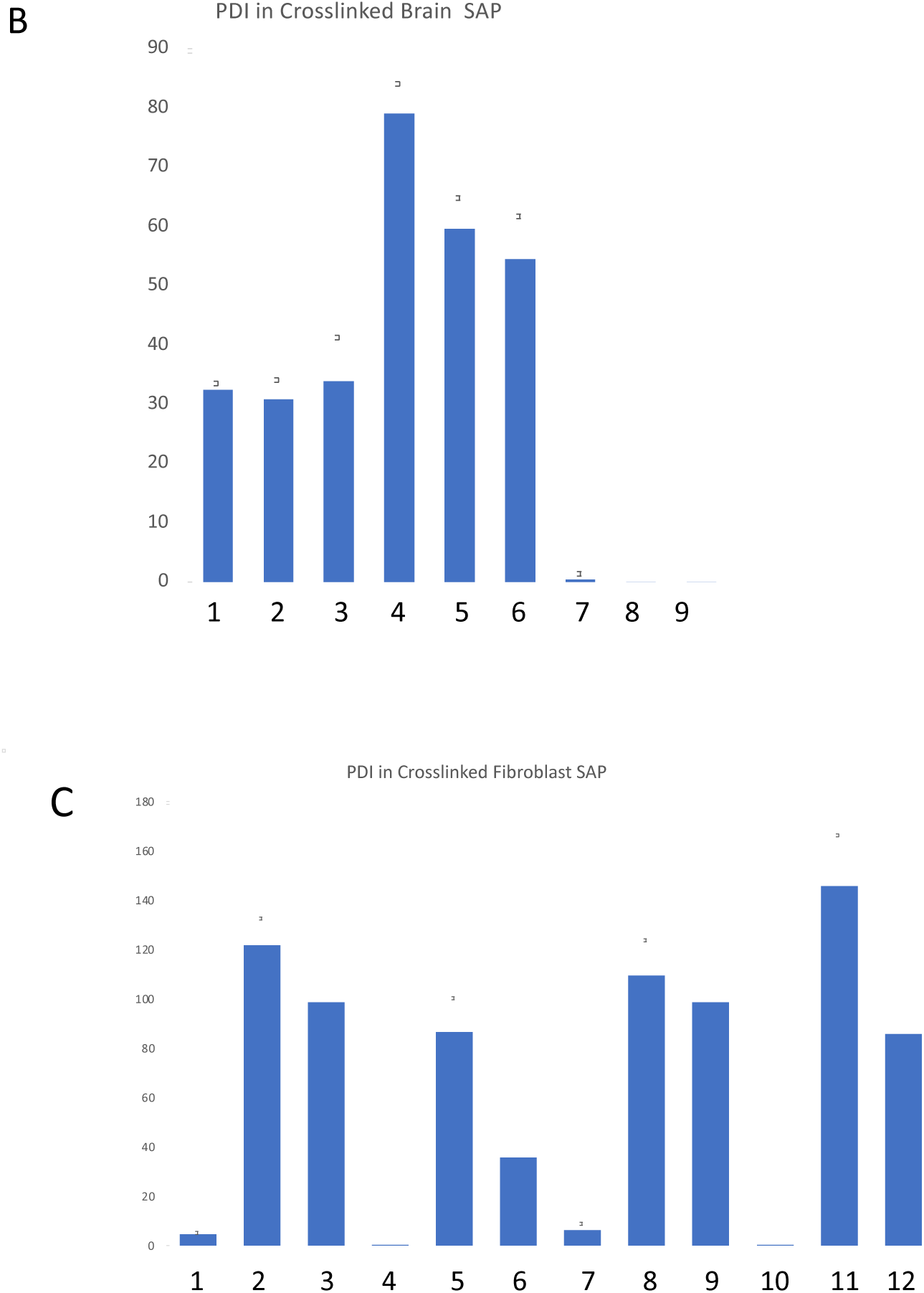
Target identification in mouse brain and PDF samples. Figure 5A shows MSMS analysis of the eDRAC eluate from a THIQ resin (30). Healthy wildtype or sick SOD1 G93A mutant mouse brain which was incubated on the THIQ resin or a control resin (affigel lacking the attached drug), washed and eluted with free compound, with the eluates analyzed by MS-MS. A list of proteins identified, and the number of spectral counts detected for each starting material and resin is shown where conditional formatting has been applied on a red-to-green scale. Protein disulphide isomerase (PDI) is detected in both drug resin eluates, but not the control resin eluates. Figure 5B shows the results of a photocrosslinking study in which healthy (bars 1, 4, 7), sick SODG93A (bars 2, 5, 8) or sick-THIQ-treated (bars 3, 6, 9) mouse brain was assessed with PAV-073 crosslinker under native (bars 1-3) or denatured (bars 4-6) conditions. Bars 7-9 show that addition of excess free drug during the crosslinking reaction eliminates PDI precipitation. This demonstration of competition between free drug and the photocrosslinker is an important line of evidence that the modification of structure occurring during crosslinker construction did not alter the capacity for target engagement. In Figure 5C photocrosslinking is done with healthy (bars 1-6) versus ALS PDF extracts (bars 7-12) handled in much the same way as in Figure 5B (native SAP bars 1-3 and 7-9; denatured bars 4-6 and 10-12), except that an additional control was performed in which crosslinker-biotin lacking the drug was shown not to precipitate PDI (bars 1, 4, 7, 10). Free drug competitor is shown in lanes 3, 6, 9, 12). For both 5B and 5C after crosslinking, the samples were divided in two, one of which was denatured in 1% SDS at 100°C for 3 minutes before addition of excess non-denaturing detergent (Triton-X-100) to take up the free SDS into micelles. Streptavidin beads were added and the bound protein was precipitated and analyzed by western blot. Quantitation of the protein band for PDI is shown as arbitrary density units, where error bars represent the averages of triplicate-repeated conditions.

To determine which of the proteins detected in the eDRAC eluate directly binds to the compound, a photocrosslinker compound was synthesized in which diazirine and biotin moieties were attached at the same position used previously to attach to the resin. Thus, upon exposure to UV light, a covalent bond is formed between the diazirine moiety of the compound and the nearest neighbor protein. Under native conditions (in the presence of metabolic energy) the full complex is isolated. However upon denaturation, followed by streptavidin precipitation (SAP), only the nearest neighbor drug-binding protein, to which an irreversible covalent crosslink has been achieved, is recovered and is identifiable by western blotting.

Compound PAV-073 was chosen as the analog to be used as a photocrosslinker for the THIQ series because its activity is selective to ALS, having largely lost efficacy against infectious HIV (see **Supplemental Figure 1** for activity of PAV-073 and **Supplemental Figure 4** for synthetic scheme of PAV-073 crosslinker). Crosslinking and SAP from mouse brain extract (brains from the SODG93A efficacy study, including wildtype, transgenic mutant, and transgenic mutant/compound treated animals) identified protein disulphide isomerase (PDI) as a direct target. Western blot analysis showed a protein band for PDI present in both native and denatured samples when crosslinked with PAV-073 but not the negative control (see **Figure 5B**). Furthermore, the PDI band was diminished with presaturation, where the free PAV-073 was added to the sample before crosslinking to compete out binding-sites (see **Figure 5B**).

The crosslinking experiment was repeated using cellular extract prepared from a sporadic ALS patient (#51) and healthy control (#27) fibroblasts (see **Figure 5C**). For both PDF and brain samples, PDI was present as a target in both healthy and ALS conditions in eDRAC and crosslinking (See **Figures 5A-C**).

## Discussion

The protein assembly modulator THIQ chemotype which is shown here active in a diversity of ALS cellular and animal models, appears to work for both familial and sporadic cases based on data shown here, generated in ALS patient-derived fibroblasts and transgenic worms, flies, and mice. Activity of hit compounds appears to normalize an array of surrogates for ALS pathology including elimination of stress-induced TDP-43 aggregates in stress granules, repair of TDP-43 mis-localization, reversal of paralysis, reversal of neurodegenerative markers, normalization of weight, and increased lifespan. Compounds appeared active on models for VCP, TDP-43, C9orf 72, and SODG93A mutations. This is consistent with an expectation that protein assembly modulation is an upstream manipulation that serves to integrate multiple biochemical pathways and thus is therapeutic for a wide range of downstream defects.

The premise in support of our unconventional approach to drug discovery and our pivot from focusing on viral to nonviral diseases once hit compounds were identified, was that viruses have used deep evolutionary time and natural selection to find the most efficient ways to take over our cells and prevent activation of host defensive measures. Thus viral targets discovered via host-viral interactome pathways likely represent weak links of human biology at risk for diseases involving departures from homeostasis-including those not caused by viruses. Experimental data appears to support this hypothesis as assembly modulator compounds have shown distinctive activity against viral disease(36), proliferative disease(35), and for neurodegenerative disease both for ALS as presented here, and previously for Alzheimer’s Disease (47).

Our findings should not be misunderstood as suggesting that viruses are causative of the neurodegenerative diseases including ALS with which they are associated. Rather our data suggest a shared molecular consequence of both viral infection and ALS: disruption of homeostasis. The data presented here suggests that this occurs by a specific molecular mechanism that involves critical components of protein assembly that can be manipulated with allosteric site-targeted protein assembly modulator drugs, to therapeutic advantage. In the case of viruses manipulation of protein assembly blocks viral capsid formation and restores the normal function of a repurposed host multi-protein complex (36). In the case of ALS a related but distinct multi-protein complex is targeted, as shown here. The precise similarities, differences, and relationship of the viral and ALS targets, and the relationship of healthy to aberrant forms occurring in disease, remains to be further studied. However the therapeutic consequences of these manipulations as shown here, suggests this to be a productive path for future effort.

We have observed that, for multiple areas of disease and multiple classes of chemical compounds, protein assembly modulators target dynamic multi-protein complexes via a protein implicated in allosteric regulation. One chemotype series active against all respiratory viruses accomplishes this by targeting 14-3-3 proteins (36,48). Another chemotype active against cancer targets KAP1 (35,49). The Alzheimer’s Disease assembly modulator targets MIF (47).The ALS/HIV active chemotype described in this paper targets PDI, a protein also implicated in allosteric regulation and in cleavage of disulfide bonds (50–53). Variants of PDI with single nucleotide polymorphisms and redistribution of PDI within different regions of the cell, including endoplasmic reticulum sub-compartments, are implicated in the literature as correlating with ALS symptoms(54–56). Furthermore, expression of PDI has been observed to protect mice against neurodegeneration in the SOD1G93A model as well as multiple cellular models of ALS (55,57–59). The PDI detected in the compound’s target by eDRAC of both healthy and ALS patient fibroblast cells, appears to be interacting with other proteins in a similarly energy-dependent manner as the other assembly modulator compounds described (35,36). As for the other assembly modulators, the protein composition of the multi-protein complex drug target differs between healthy and sick cells (Figure 5A), although we do not yet have compelling evidence, as observed for other assembly modulators(36), for normalization of a disease-associated target. Possible explanations include the need to continue to drive SAR further towards disease-selectivity, that the specific allosteric site targeted is shared by both healthy and sick individuals with disease-specific features not reflected in the analysis to date.

We hypothesize that the unique properties observed in multiple distinct chemical classes of assembly modulators may be attributable to their having a mechanism of action that works via allosteric regulation. Targeting allosteric sites instead of active sites may be an effective strategy to selectively inhibit activity of the subsets of proteins which have been dysregulated in a diseased state, without targeting the forms in the healthy state. Modulation of the multi-protein complex that catalyzes protein assembly may provide a means of restoring homeostasis, rather than simply blocking a disease-associated form, without restoration of homeostasis.

Our use of viral assembly as a surrogate for discovery of small molecule protein assembly modulators of relevant allosteric sites for the restoration of homeostasis has proven productive. In part, this may be because it is not constrained by the limitations of current technology. Identifying allosteric sites, which affect activity of a small subset of a given gene product cannot be easily achieved through genetic manipulation, given the evidence for protein “moonlighting”(60,61). Of necessity, genetic manipulation affects all of the diversity of forms of a particular gene product, irrespective of the different functional roles they may play. Since only one of these functions may be responsible for the disease, targeting all forms and their functions, may obscure therapeutic effect, e.g. through off-target liabilities. Finding a small molecule that selectively targets only the relevant subset of protein-protein interactions provides a path to retain broad activity while avoiding toxicity, as demonstrated for other protein assembly modulator compounds(35,36), and suggested for ALS from the data presented here.

Efficacy of the same protein assembly modulator small molecule in both familial and sporadic ALS supports the hypothesis that these compounds act on an upstream regulatory mechanism rather than on downstream consequences of disease (manifest as protein aggregates). Thus one therapeutic small molecule is broadly applicable despite the heterogeneity of ALS. As an analogy, when car crashes frequently occur at a busy intersection, upstream solutions, like installing a stop sign or fixing a broken traffic light, are more effective than a focus on downstream consequences, such as sending more tow trucks to remove roadside wreckage. Sending tow trucks to remove the wreckage will improve traffic in the short term, but if the underlying defect which led to the crash is not addressed— it is only a matter of time before it happens again. Targeting the aggregates themselves is akin to eliminating wreckage solely by sending tow trucks following a crash. Allosteric modulation on the other hand, is a way to prevent wreckage by installing a stop sign to regulate traffic from a distance(62). Figure 6 illustrates the distinction between conventional thinking with regards to protein aggregate-associated diseases including ALS, and the hypotheses formulated and tested here.

**Figure 6.**
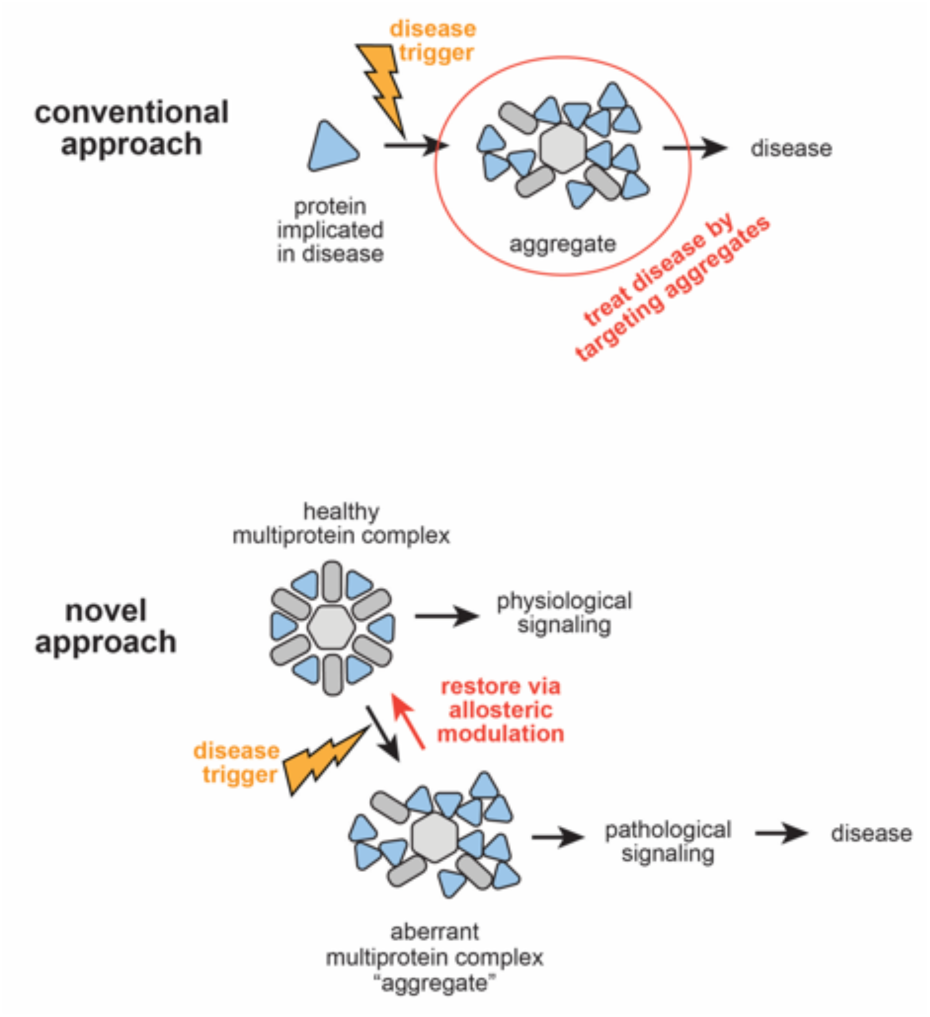
Protein assembly modulation drug action. Hypothesized mechanism of protein assembly modulator drug action for protein aggregation diseases including ALS, based on the data. Top: conventional approaches aimed at removal of aggregates. Bottom: Protein assembly modulation approach to preventing aggregate formation. By targeting an allosteric site to prevent aggregate formation, rather than targeting aggregates after they have formed, the outcome is akin to installing a stop sign to prevent accidents at a busy intersection, rather than sending tow trucks to remove wreckage once an accident has occurred.

Another implication of the findings presented here is that, though the targets of the viral and related non-viral diseases are similar, they are nevertheless sufficiently distinctive that their activity can be separated through SAR advancement. We observed early compounds in the THIQ series show potency in models for both ALS and HIV. However, the advanced ALS-active compound PAV-073, demonstrated potent activity in the worm model for ALS while showing substantial loss of activity against infectious HIV in cell culture. Not shown, a different advanced subseries is HIV-selective, without activity in an ALS model. This suggests that there are two separate targets (one relevant for ALS, one for HIV) for this chemical series. Driving the SAR towards selectivity will likely diminish liability for off-target toxicity and illuminate the molecular mechanisms underlying each disease state in ways that have been heretofore inaccessible.

## Supplemental Figures

**Supplemental Figure 1.**
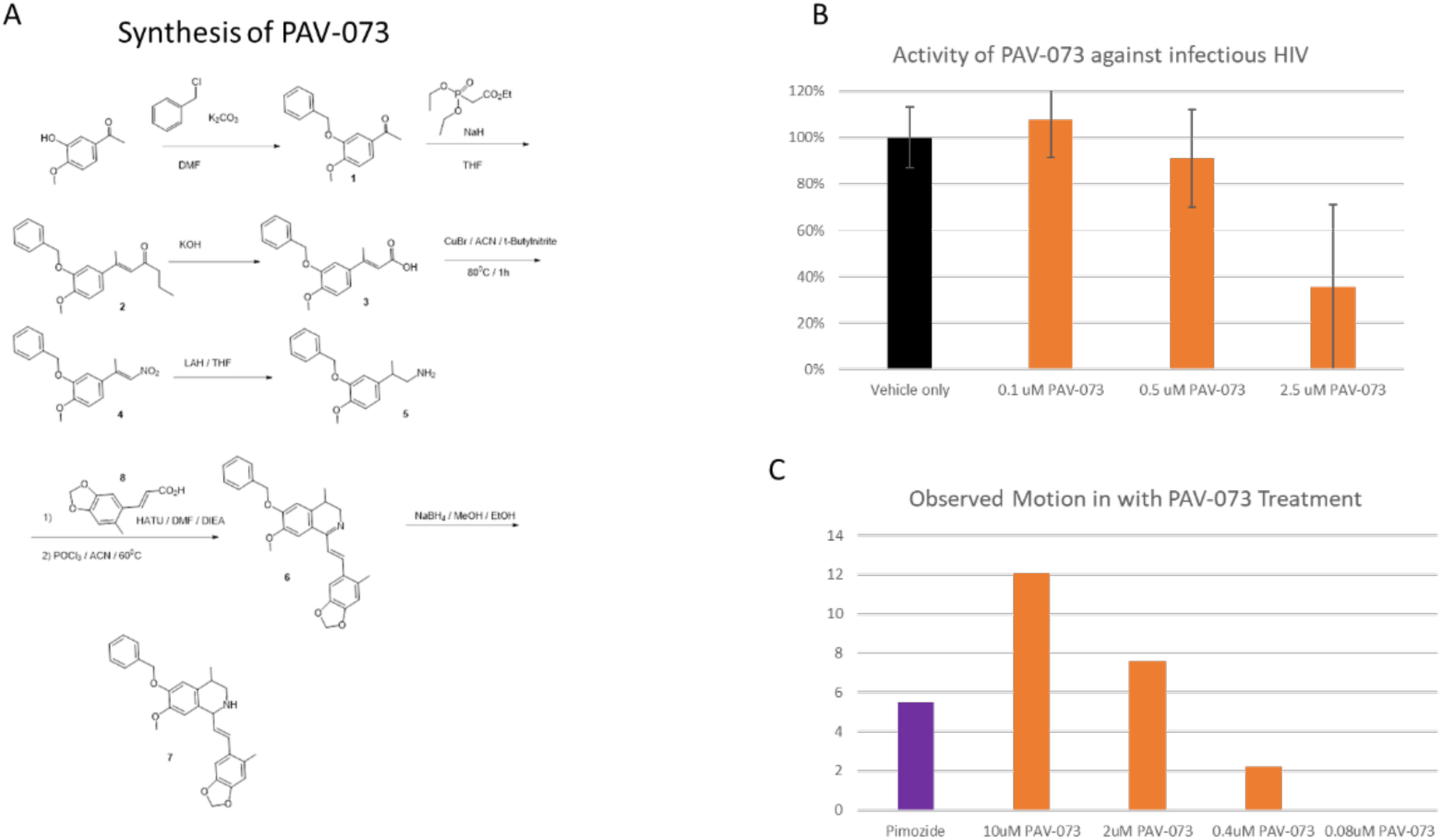
Synthesis and activity of PAV-073. **Supplemental Figure 1A** shows the synthetic scheme for PAV-073. **Supplemental Figure 1B** shows activity of PAV-073 against infectious HIV. MT-2 cells were infected with NL4-3 Rluc HIV and treated with PAV-073 for four days. Averages and standard deviation of viral titer observed with triplicate repeated dose-titrations of PAV-073 are shown as a percentage of the titer observed in DMSO-treated cells. **Supplemental Figure 1C** shows activity of PAV-073 relative to pimozide (40uM) in ameliorating the condition of transgenic *C. elegans* expressing the human TDP-43 A315T mutation. Nematodes were age-matched and grown on standard nematode grown media plates until day 1 of adulthood at which point they were collected and placed in 96 well plates (50-70 animals per well) and treated with compound or control. Animal movement was then tracked for 30 minutes using WMicroTracker ONE.

**Supplemental Figure 2.**
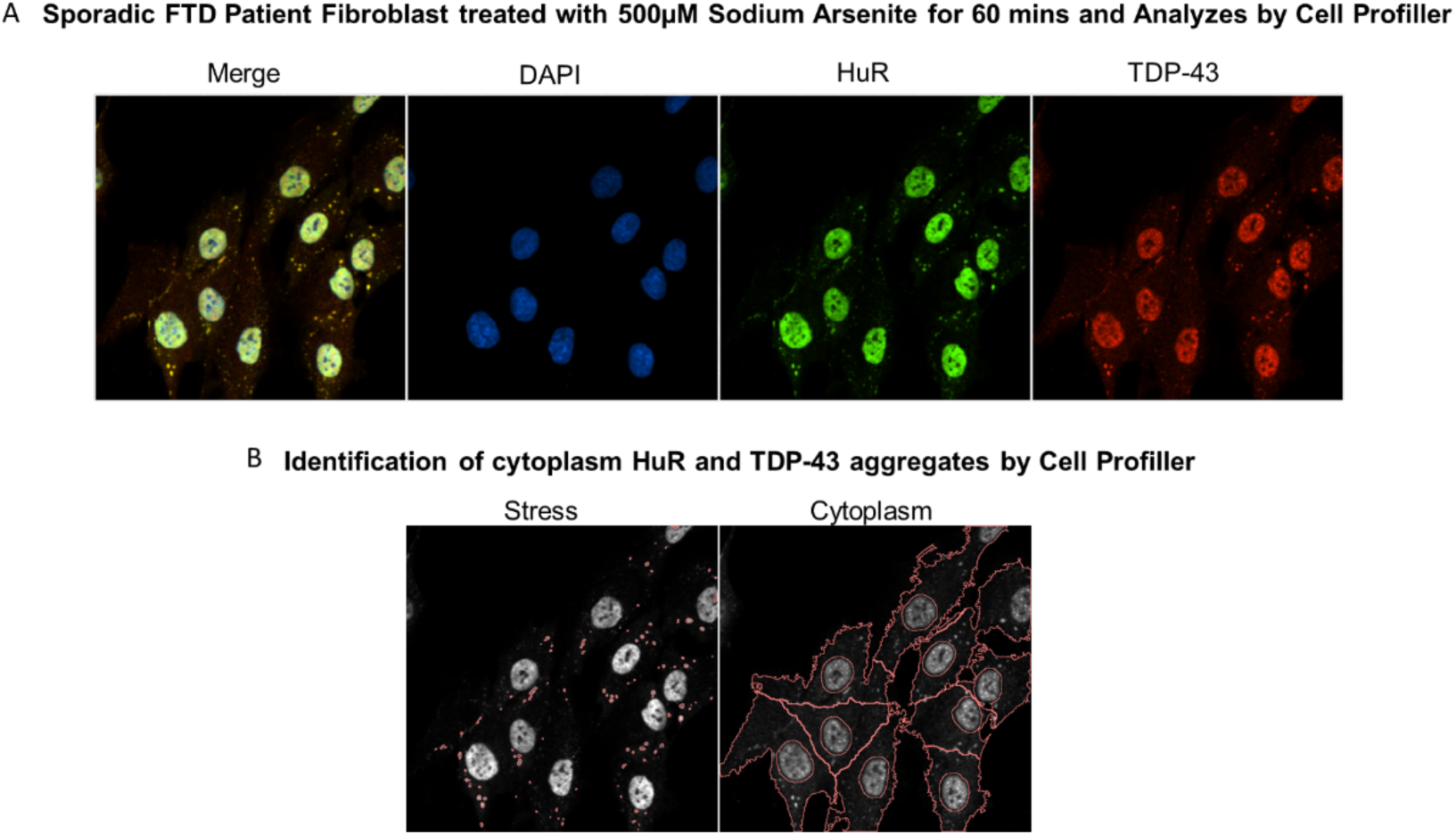
Stress granule induction and quantitation. **Supplemental Figure 2A** shows immunostain for DAPI, HuR, and TDP-43 in PDFs treated with 500uM sodium arsenite for one hour. **Figure 2B** shows how cell profiler imaging was used to identify the number of TDP-43 positive HuR aggregates per cell.

**Supplemental Figure 3.**
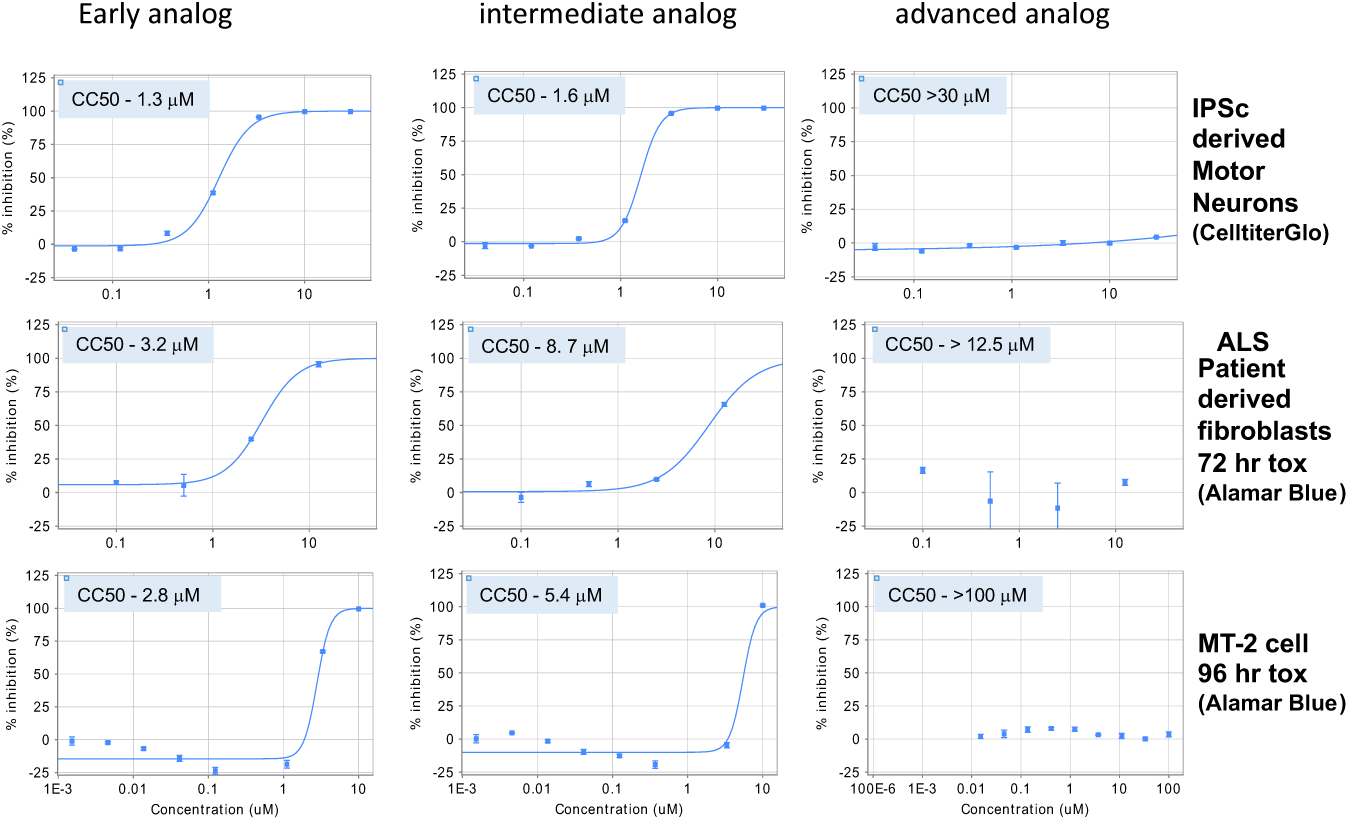
Moderation of toxicity with THIQ lead series advancement. Compounds of comparable potency but differences in toxicity were identified from the lead series and assessed in transformed cells, ALS PDFs, and iPSC-derived motor neurons. The lead series progression to diminished toxicity is confirmed in all cell lines using both cell TiterGlo and Alamar Blue toxicity assays.

**Supplemental Figure 4.**
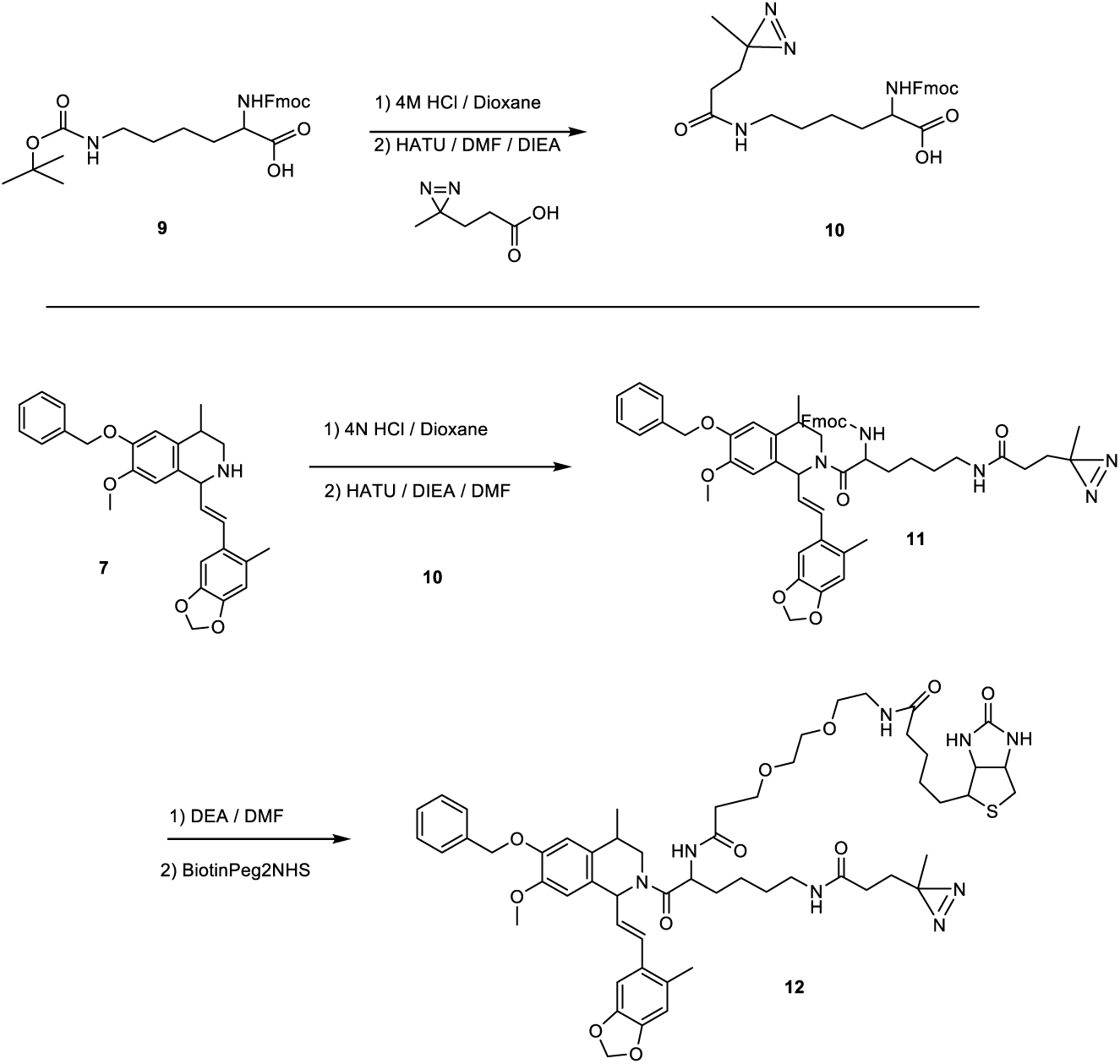
Synthetic scheme for PAV-073 photocrosslinker analog. **Supplemental Figure 3** shows the synthetic scheme for the photocrosslinker analog of PAV-073 used to identify PDI as the direct drug binding protein (see **Figure 5**).

## Materials and Methods

### Lead contact and Materials Availability

Further information and requests for resources and reagents should be directed to and will be fulfilled by the Lead Contact Vishwanath R. Lingappa.

Use of unique compound PAV-073 and its stable derivatives may be available upon request by the Lead Contact if sought for experimental purposes under a valid completed Materials Transfer Agreement.

### Chemical Synthesis (see Supplemental Figure 1 and Supplemental Figure 3)

#### Synthesis of PAV-073

A mixture of 3-hydroxy-4-methoxyacetophenone (16.6 g, 100 mmol), benzyl chloride (13.8 mL,120 mmol), and anhydrous K2CO3 (20.7 g, 150 mmol) in DMF (100 mL) was heated at refluxed for 5 h. The reaction mixture was concentrated to dryness, the residue was redissolved in EtOAc (100 mL) and then washed with 5%aqueous NaOH (3x30 mL). The organic layer was washed with brine (2x10 mL) and H2O (2x30 mL), dried (Na2SO4) and evaporated to a residue, which was purified by flash chromatography to provide 1 (22.9 g, 90%). MS (m/z):257 [M+H].

NaH (60 wt % in mineral oil, 1.95 g, 48.5 mmol) was suspended in THF (100 mL) and cooled to 0° C. Triethylphosphonoacetate (9.6 mL, 48.5 mmol) was added dropwise and the reaction mixture was stirred at 0° C. for 30 min. Then 3-benzyloxy-4-methoxy-acetophenone (1) (6.2 g, 24.2 mmol) was dissolved in THF (0.1 ml/mmol) and added to the reaction mixture. The cooling bath was removed and the mixture was stirred at 50° C. until full conversion was detected (TLC). The reaction mixture was quenched by slow addition of H2O (2 ml/mmol ketone), extracted with t-butyl methyl ether (3x3 ml/mmol) and the combined organic layers were dried (Na2SO4) and evaporated to give a residue, which was purified by flash column chromatography to provide compound 2 (6.4 g, 81%). MS (m/z): 327 [M+H].

A mixture of ethyl ester (2) (6.4 g, 19.5 mmol) and alcoholic potassium hydroxide (4.0 g, 71 mmol KOH/ 100 mL EtOH) was stirred at room temperature for 12 h. The solution was then concentrated to give a residue, which was purified by flash column chromatography on silica gel to provide 3 (5.6 g, 96%). MS (m/z): 299 [M+H].

A suspension of 3-(3-benzyloxy-4-methoxyphenyl)-2-butenoic acid (3) (5.6 g, 18.8 mmol), CuBr (270mg, 1.9 mmol) and tertiary butyl nitrite (8.9 mL, 37.6 mmol) in acetonitrile (50 mL) was stirred at 80° C. for 18 h. Reaction completion was monitored by TLC. After completion, the reaction mixture was cooled to room temperature, solvent was removed under reduced pressure and the crude product was purified by flash chromatography to yield compound 4 (3.9 g, 70%). MS (m/z): 300 [M+H].

To a solution of 3-(3-benzyloxy-4-methoxyphenyl)-l-nitro-2-butene (4) (3.9 g, 13.2 mmol) in 40 mL of anhydrous THF under argon was slowly added a 2.0 M solution of LiAlH4 in THF (40 mL, 80 mmol) and the reaction mixture was heated at refluxed for 2 h. The reaction mixture was cooled and excess reagent was quenched by dropwise addition of H2O and 15% aqueous NaOH. The reaction mixture was extracted with CH2Cl2 (3x30 mL) and the combined organic layers were treated with 5% aqueous HC1. The aqueous acid layer was then basified (5% aqueous NH4OH, pH 9) and extracted with CH2Cl2. The organic solution was washed with brine (2x30 mL) and H2O (2x30 mL), dried (Na2SO4) and evaporated to give compound (5) (2.3 g, 63%). MS (m/z): 272 [M+H].

To the stirred solution of (E)-3-(6-methyl-l,3-benzodioxol-5-yl)prop-2-enoic acid (8) [see synthesis below] (140 mg, 0.68 mmol)and 2-(3-benzyloxy-4-methoxy-phenyl)propylamine (5) (185 mg, 0.68 mmol) in DMF (2 mL) was added HATU (310 mg, 0.82 mmol) and diisopropylethylamine (351 mg, 0.473 mL, 15.0 mmol). The reaction mixture was stirred at room temperature for 1 h, diluted with EtOAc (50 mL), washed with 10% citric acid, saturated aqueous solution of NaHCO3, dried (Na2SO4), filtered and evaporated to give a residue, which was purified by flash chromatography (ethyl acetate/hexanes) to provide compound 6. Yield 11.4 mg (35% overall yield from nitrostyrene). MS (m/z): 460 [M+H].

A suspension of (E)-N-[2-(3-benzyloxy-4-methoxy-phenyl)-propyl]-3-(6-methyl-l,3-benzodioxol-5-yl)-prop-2-enamide (6) (110 mg, 0.24 mmol) in dry acetonitrile (10 mL) was heated at reflux. Then phosphorus oxychloride (400 mg, 0.24 mL, 2.6 mmol) was added drop wise and the reaction mixture was heated at reflux for an additional 1 h. The solvent and reagent were evaporated under vacuum, the organic layer was washed with water (2x10 mL). and evaporated in vacuo to give an oil, which was then dissolved in ethanol (8 mL) and sodium borohydride (9.8 mg, 0.26 mmol) was added. The reaction mixture was stirred at room temperature for 30 min and excess reagent was destroyed by dropwise addition of 2 M HC1. The reaction mixture was basified with 2 M NaOH and ethanol was removed in vacuo to give a residue, which was partitioned between water (10 mL) and chloroform (10 mL). The organic layer was washed with water (2x10 mL), dried and evaporated to give a residue, which was purified by column chromatography (dichloromethane/methanol) to give 6-nenzyloxy-7-methoxy-4-methyl-l-[(E)-2-(6-methyll, 3-benzodioxol-5-yl)-vinyl]-1,2,3,4-tetrahydroisoquinoline (7) (10 mg, 10%). MS (m/z): 444 [M+H].

#### Synthesis of PAV-073 Photocrosslinker

To 6-(tert-Butoxycarbonylamino)-2-(9H-fluoren-9-ylmethoxycarbonylamino)-hexanoic acid 9 [468mg (1mmol)]a in a 40ml screw top vial was added 4N HCl in Dioxane (3ml). The vial was sealed and gently agitated for 20min at room temperature. The mix was then rotary evaporated to dryness and the residue placed on high vacuum overnight.

The dried residue was taken up into 4ml of DMF (anhydrous) and then sequentially treated with 3-(3-Methyldiazirin-3-yl)-propanoic acid [128mg (1mmol)]b, and DIEA [695ul (4mmol)]. With rapid stirring, under Argon atmosphere, was added dropwise HATU [380mg (1mmol)] dissolved in 1ml of DMF. After stirring for 30 min the mixture was quenched with 10ml of sat. NH4Cl solution and then extracted 2 x with 10ml of EtOAc.

The combined organic extracts were washed once with sat. NaCl, dried (Mg2SO4) and then rotary evaporated to dryness. The residue was purified by flash chromatography, using a gradient of Ethyl acetate and Hexane, affording 10 (293mg) in 61% yield.

To 6-benzyloxy-7-methoxy-4-methyl-1-[(E)-2-(6-methyl-1,3-benzodioxol-5-yl)vinyl]-1,2,3,4-tetrahydroisoquinoline 7 [15mg (0.03 mmol)]c in a 40ml screw top vial was added 4N HCl in Dioxane (0.5ml). The vial was sealed and gently agitated for 20min at room temperature. The mix was then rotary evaporated to dryness and the residue placed on high vacuum overnight.

The dried residue was taken up into 1ml of DMF (anhydrous) and then sequentially treated with 10 [14.5mg (0.03mmol)], and DIEA [32ul (0.18mmol)]. With rapid stirring, under Argon atmosphere, was added dropwise HATU [14.6mg (0.038mmol)] dissolved in 300ul of DMF. After stirring for 30 min the mixture was quenched with 5ml of sat. NH4Cl solution and then extracted 2 x with 5ml of EtOAc.

The combined organic extracts were washed once with sat. NaCl, dried (Mg2SO4) and then rotary evaporated to dryness. The residue was purified by flash chromatography, using a gradient of Ethyl acetate and Hexane, affording 11 (28mg) in quant. yield.

To 11 [28mg (0.03 mmol)] in a 40ml screw top vial was added 50/50 Diethylamine / DMF (0.5ml). The vial was sealed and gently agitated for 60min at room temperature. The mix was then rotary evaporated to dryness and the residue placed on high vacuum overnight. The residue was triturated 2 x with 3ml of Hexane to remove the Dibenzofulvene amine adduct. The residue was again briefly placed on high vacuum to remove traces of Hexane. The dried residue was taken up into 1ml of DMF (anhydrous) and then treated with Biotin-PEG2-NHS [15mg (0.03mmol)]d, and DIEA [16ul (0.09mmol)] and then purged with Argon. After stirring overnight at room temperature, the mixture was rotary evaporated to dryness. The residue was purified by reverse phase prep chromatography, using a gradient of 0.1% TFA water and Acetonitrile, affording 12 (26mg) in 80% yield. Purity of all compounds were confirmed via LCMS.

## Method and Analysis Details

### In vitro studies

#### Stress granule aggregation in SH-SY5Y cells overexpressing TDP-43 SH-SY5Y cells

A TDP-43 cellular model was successfully generated by establishing stable cell-lines over-expressing wild-type TDP-43 and M337V mutant TDP-43 and showing the display of stress granules, following arsenite treatment. SH-SY5Y tet-on TDP-43 partial 3′-UTR (wt or M337V), 50.000 cells (DMEM/F12 + 1 µg/mL doxycycline) incubated on cover slips overnight. Cells were treated sodium arsenite at a final concentration of 250 µM and incubated for 90 min. Cells were washed with 500 µL PBS. Cells were fixed with 4% PFA in PBS pH 7.4 for 15 min at RT. Cells were washed with 500 µL PBS. Permeabilize/ block cells with 5% milk powder, 1% BSA and 0.5 % saponin in PBS for 45 min @ RT. Add respective {TDP-43 C-terminal domain Antibody (1:1000; Purchased from Proteintech), HuR (1:500; Purchased from Santa Cruz} in 1% PBS and 0.5% saponine in PBS and incubated overnight at 4°C. Cells were washed 3 x with 500 µL PBS and incubated with secondary antibody at a dilution of 1:1000 AlexaFluor a-rabbit 594 (highly cross-adsorbed) (Thermo Fisher) or AlexaFluor a-mouse 488 (highly cross-adsorbed) (Thermo Fisher). Cells were washed 3x with 500 µL PBS, then cells were washed with 500 µL dH2O. ProLong Gold with DAPI embedding medium was used to fix cells on a glass slide. Image collection was done on a Zeiss AxioImager 2 equipped with an Apotome using the following filtersets. For the automated image analysis, each raw grayscale channel image was saved and analyzed independently. Each image set was calibrated using the TDP-43 induced, sodium arsenite treated condition as a reference for exposure time of the different channels.

#### High-Content Imaging of Endogenous TDP-43 Stress Granule (SG)

On day 1, seed 20,000 patient derived fibroblasts per well in a 24 well glass bottom plate or 6000 cells per well in a 96 well plate. On day 2, sonicate compounds for 10 mins at 37oC before use. Add compounds at the desired final concentration in fresh media to the respective wells. Add equivalent amount of DMSO (LC-MS grade) to control wells. On Day 3, add sodium arsenite treatment-Add sodium arsenite at a final concentration of 500uM. Incubate at 37oC for 60 mins. Wash 1X with PBS and fix cells with 4% para formaldehyde (in PBS, prepared freshly, methanol free) for 15 mins at room temperature.

Wash 3X with PBS. Permeabilization and Blocking-Add 0.1% Triton-X for 10 mins for permeabilization followed by 1 hour of blocking in 1% BSA. Immunostaining-Add the following primary antibodies in 1% BSA (in PBS) and incubate it overnight at 4oC. Rabbit polyclonal TDP-43 C-terminal antibody (Proteintech 12892-1-AP)-1:450; mouse monoclonal HuR antibody (Santa Cruz sc-5261)-1:500. On day 4, wash 3X with PBST (PBS + 0.1% Tween). The following secondary antibodies from Thermofisher Scientific (1:500) in 1% BSA (in PBS) and keep it in dark for 1-2 hours at room temperature.

Alexa 594 anti-rabbit (highly cross-adsorbed); Alexa 488 anti-mouse (highly cross-adsorbed); Wash 3X with PBST in dark. Add DAPI in PBS for nuclear staining. Imaging and Image analysis is done as explained below. In brief, the immuno-stained cells were imaged with Nikon Ti inverted fluorescence microscope having CSU-22 spinning disk confocal and EMCCD camera. Plan Apo objectives and NIS-Elements AR software were used for image acquisition. At least 30-50 images per well is taken.

#### Nucleocytoplasmic assay in FTD patient fibroblasts cells

Skin-derived fibroblasts cells from a sporadic Frontotemporal Degeneration (FTD) and ALS affected individual acquired from the National Institute of Neurological Disorders and Stroke were grow in HyClone DMEM High Glucose (GE Healthcare Life Sciences) supplemented with 15% FBS and 1% NEAA (Non-Essential Amino Acids), at 37°C in an humidified atmosphere of 5% CO2. On Day 1, seed 600 cells per well in a 96 well glass bottom plate or 1200 cells per well in a 24 well glass bottom plate. Incubate for 4 days, at 37°C in an humidified atmosphere of 5% CO2. On day 5, sonicate compounds for 10 mins at 37oC before use. Add compounds at the desired final concentration in fresh media to the respective wells. Add equivalent amount of DMSO (LC-MS grade) to control wells. Incubate for 4 days, at 37°C in a humidified atmosphere of 5% CO2. On day 9, wash 2X with PBS. To fix add 4% para formaldehyde (in PBS, prepared freshly, methanol free) for 15 mins at room temperature. Wash 3X with PBS. Blocking and permeabilization-Add 1% BSA + 1% saponin (prepared in PBS) for 1 hour.

Immunostaining-Add the following primary antibodies in 1% BSA (in PBS) and incubate it overnight at 4oC. Rabbit polyclonal TDP-43 C-terminal antibody (Proteintech 12892-1-AP)-1:350; mouse monoclonal HuR antibody (Santa Cruz sc-5261)-1:500. On day 10, wash 3X with PBST (PBS + 0.1% Tween). Add the following secondary antibodies from Thermofisher Scientific (1:500) in 1% BSA (in PBS) and keep it in dark for 1-2 hours at room temperature. Alexa 594 anti-rabbit (highly cross-adsorbed). Alexa 488 anti-mouse (highly cross-adsorbed). Wash 3X with PBST in dark. Add DAPI in PBS for nuclear staining. The immuno-stained cells are imaged with Nikon Ti inverted fluorescence microscope having CSU-22 spinning disk confocal and EMCCD camera. Plan Apo 20x/0.75 objective and NIS-Elements AR software were used for image acquisition. At least 15 images per well are taken. The exposure times for TDP-43 and HuR must remain constant across one experiment. Each image acquired (in .nd format) is exported into three individual channel images (for DAPI, TDP-43, HuR) in .tiff format. The images are analyzed by the open source image analysis software Cell Profiller. The DAPI image is used to count the total number of cells. The TDP-43 and HuR images are used to count the number of cells containing TDP-43 and/or HuR nuclear staining.

#### Automated Image Analysis and Machine Learning Tools

The immuno-stained cells were imaged with Nikon Ti inverted fluorescence microscope having CSU-22 spinning disk confocal and EMCCD camera. Plan Apo objectives and NIS-Elements AR software were used for image acquisition. At least 30-50 images per well is taken. The exposure times for TDP-43 and HuR must remain constant across one experiment. Each image acquired (in .nd format) is exported into three individual channel images (for DAPI, TDP-43, HuR) in .tiff format. The images were analyzed by the open source image analysis software CellProfiler 2.1.1 (offered by Broad Institute of Harvard and MIT-www.cellprofiler.org). This software contains various modules which can be used to analyze images in different ways. A Cell Profiler pipeline from a few of these modules was established to analyze our images in order to quantify the number of cytoplasmic TDP-43 positive HuR stress granules. The outline of the Cell Profiler pipeline: The three channel images are loaded and named DAPI, TDP-43 or HuR. Identify primary objects-The DAPI image is used to identify the nucleus as an object. Identify secondary objects-The nucleus is used to identify the cell boundary in the TDP-43 image by signal propagation. Identify tertiary objects-Based on the nucleus and the cell boundary, cytoplasm is identified as an object. Mask images-Using the cytoplasm object, the TDP-43 and HuR images are masked such that only cytoplasmic signal will remain. Enhance features: Enhance the signal from TDP-43 and HuR aggregates in cytoplasm for efficient identification of the aggregates. Identify primary objects-The TDP-43 aggregates in the cytoplasm were identified from the TDP-43 image and HuR aggregates were identified from the HuR image. Relate objects-This module enables the calculation of the number of TDP-43 aggregates which has HuR and vice versa. Export to spreadsheet-This module exports all data into Excel sheets. The final data has to be curated from the Excel sheets generated by Cell Profiler. The outlines for the different objects (nucleus, cytoplasm, aggregates) must be saved to cross-check the proper identification of objects once the analysis is done. At the beginning of analysis of an experiment, few images (from DMSO wells) must be used as training set for Cell Profiler. Based on the training set, the pipeline must be optimized with respect to intensity threshold, algorithm and size parameters for correct identification of primary and secondary objects (nucleus, cytoplasm, aggregates). It is extremely important to optimize this pipeline for every experiment. The pipeline must remain constant with respect to aggregate identification for the analysis of all the images from the same experiment. Finally, we calculate the number of TDP-43 positive HuR stress granules from the Excel sheets generated at the end of Cell Profiler.

### HIV infectious virus assay

MT-2 cells were preseeded in 96-well plates in 100 ul of complete RPMI. Multiple concentrations of PAV-951 were serially diluted in DMSO then into an infection media prepared by diluting NL4-3 Rluc virus stock to 400 IU/100 ul with complete RPMI, which was transferred onto the MT-2 cells with a final MOI of 0.02 and final DMSO concentration of 1% in infected places. One well received DMSO only, instead of PAV-951, and one well received medium only for normalization and background collection. Cells were incubated at 37° C for 96 hours. 100ul of medium was removed and discarded and 10 ul of 15 uM EnduRen luciferase substrate was added to each well, followed by incubation for 1.5 hours at 37° C. Plates were read on a luminescence plate reader. Bioluminescence intensity was read on a Synergy H1 BioTek plate reader. Averages and standard deviation for viral titer observed under different treatment conditions were calculated in Microsoft Excel and graphed as the percent inhibition in PAV-951 treated cells compared to untreated cells.

### Drug Resin affinity chromatography

Mouse brains from wildtype or SOD1 mutant animals were homogenized in cold phosphate buffered saline (PBS) (10mM sodium phosphate, 150 mM sodium chloride pH 7.4), then spun at 1,000 rpm for 10 minutes until pelleted. The PBS was decanted and the pellet resuspended in a low salt buffer (10mM HEPES pH 7.6, 10mM NaCl, 1mM MgAc with 0.35% Tritonx100) then centrifuged at 10,000 rpm for 10 minutes at 4°C. The post-mitochondrial supernatant was removed and adjusted to a concentration of approximately 10 mg/ml and equilibrated in a physiologic column buffer (50 mM Hepes ph 7.6, 100 mM KAc, 6 mM MgAc, 1 mM EDTA, 4mM TGA). In some conditions, the extract was supplemented with an energy cocktail (to a final concentration of 1mM rATP, 1mM rGTP, 1mM rCTP, 1mM rUTP, and 5 ug/mL creatine kinase). 30 ul or 230 ul of extract was then incubated for one hour at either 4° C or 22° C degrees on 30 ul of affigel resin coupled to THIQ compound or a 4% agarose matrix (control). The input material was collected and the resin was then washed with 3 ml column buffer. The resins were eluted for 2 hours then overnight at 22°C then 4°C in 100ul column buffer containing 100uM of the cognate compound. Eluates were run on western blot or sent for mass spectrometry for analysis.

### Chemical photocrosslinking

Extract from mouse brain and PDFs grown in minimum essential media were prepared as above then adjusted to a protein concentration of approximately 3 mg/ml in column buffer containing 0.01% triton. 1% DMSO or 100uM PAV-073 was added to 6ul of extract, then 3uM of PAV-073 photocrosslinker or a negative control crosslinker (comprising of the biotin and diazirine moieties without compound) were added. The extract was incubated for 20 minutes then exposed to UV at 365nM wavelength for 10 minutes then left on ice for one hour. After crosslinking, samples were divided in two 20 ul aliquots and one set was denatured by adding 20 uL of column buffer 4ul of 10% SDS, 0.5 ul 1M DTT, and boiling for 5 minutes. Both native and denatured aliquots were then diluted in 800 ul column buffer containing 0.1% triton. 5 ul of magnetic streptavidin beads (Pierce) were added to all samples and mixed for one hour at room temperature to capture all biotinylated proteins and co-associated proteins. Samples were placed on a magnetic rack to hold the beads in placed and washed three times with 800 ul of column buffer containing 0.1% triton. After washing, beads were resuspended in 80 ul of gel loading buffer containing SDS and analyzed by western blot or blot for affinity purified streptavidin. Samples were analyzed by western blot.

### Western blotting

SDS/PAGE gels were transferred in Towbin buffer (25mM Tris, 192mM glycine, 20% w/v methanol) to polyvinylidene fluoride membrane, blocked in 1% bovine serum albumin (BSA) in PBS, incubated overnight at 4°C in a 1:1,000 dilution of 100ug/mL affinity-purified primary IGG to PDI in 1% BSA in PBS containing 0.1% Tween-20 (PBST). Membranes were then washed twice in PBST and incubated for two hours at room temperature in a 1:5000 dilution of secondary anti-rabbit or anti-mouse antibody coupled to alkaline phosphatase in PBST. Membranes were washed two more times in PBST then incubated in a developer solution prepared from 100 uL of 7.5 mg/mL 5-bromo-4-chloro-3-indolyl phosphate dissolved in 60% dimethyl formamide (DMF) in water and 100ul of 15 mg/ml nitro blue tetrazolium dissolved in 70% DMF in water, adjusted to 50mL with 0.1 Tris (pH 9.5) and 0.1 mM magnesium chloride. Membranes were scanned and the integrated density of protein band was measured on ImageJ. Averages and the standard deviation between repeated experiments were calculated and plotted on Microsoft Excel.

### Tandem mass spectrometry

Samples were processed by SDS PAGE using a 10% Bis-tris NuPAGE gel with the 2-(N-morpholino)ethanesulfonic acid buffer system. The mobility region was excised and washed with 25 mM ammonium bicarbonate followed by 15mM acetonitrile. Samples were reduced with 10 mM dithoithreitol and 60° C followed by alkylation with 5o mM iodoacetamide at room temperature. Samples were then digested with trypsin (Promega) overnight (18 hours) at 37° C then quenched with formic acid and desalted using an Empore SD plate. Half of each digested sample was analyzed by LC-MS/MS with a Waters NanoAcquity HPLC system interfaced to a ThermoFisher Q Exactive. Peptides were loaded on a trapping column and eluted over a 75 uM analytical column at 350 nL/min packed with Luna C18 resin (Phenomenex). The mass spectrometer was operated in a data dependent mode, with the Oribtrap operating at 60,000 FWHM and 15,000 FWHM for MS and MS/MS respectively. The fifteen most abundant ions were selected for MS/MS.

Data was searched using a local copy of Mascot (Matrix Science) with the following parameters: Enzyme: Trypsin/P; Database: SwissProt Human (conducted forward and reverse plus common contaminants); Fixed modification: Carbamidomethyl (C) Variable modifications: Oxidation (M), Acetyl (N-term), Pyro-Glu (N-term Q), Deamidation (N/Q) Mass values: Monoisotopic; Peptide Mass Tolerance: 10 ppm; Fragment Mass Tolerance: 0.02 Da; Max Missed Cleavages: 2. The data was analyzed by spectral count methods.

### In vivo studies

#### Transgenic Human TDP-43 mutant *C. elegans*

MosSCI homologous-recombination transgenesis was used to create an unc-47p::hTDP-43::unc-54utr or unc-47p::hTDP-43(mutant M337V)::unc-54utr transgenic. Transgenesis requires MOSSCI plasmid inserted with unc-47p::hTDP-43::unc-54utr or unc-47p::hTDP-43(mutant M337V)::unc-54utr. Injection mix used Standard MosSCI mix. Injections were performed into mos1 ttTi5605 background strain. Extrachromosomal array lines were isolated. Crawling transgenics screened as non-red homozgotes were verified by PCR for insertion/replacement at target locus resulting verified single copy integrated strains. Transgenic *C. elegans* expressing the human TDP-43 wild-type or mutant TDP-43 M337V animal model that mimic aspects of TDP-43 specific ALS disease pathogenesis were generated. The transgenic *C. elegans* had a single copy of the human TDP-43 gene integrated into its genome. The expression is controlled by an unc-47 promoter and hence human TDP-43 protein was specifically expressed only in the *C. elegans* motor neurons. C *elegans* studies were also performed with worms transgenic for the hTDP-43 (A315T) mutation using methods described in detail elsewhere(23,45).

#### Age-synchronizing C. elegans

Filtered deionized water is used to wash worms off of plates and into 15ml tubes which are centrifuged at 1200 rpm for 2 minutes and repeated twice. The supernatant is aspirated and 5ml of NaOH + bleach solution added. This is vortexed gently about every minute and monitored by microscope. The adults worms split open and their eggs are released. The adult worms also dissolve into the solution. Once all adult worms have dissolved, the reaction is neutralized by adding 5 ml of M9 buffer followed by three rounds of centrifugation at 2500 rpm for 2 minutes. After one wash with 10 ml of water, all but about 200-1000ul is aspirated from the 15ml tube and the remaining pellet will be re-suspended in leftover water. This are dropped onto the plates evenly, thus ensuring that the larva that hatches have enough food while they grow over the next few days. Plates will be stored at 20°C.

#### Swimming-induced Paralysis (SWIP) Assay

The age-synchronized worms are washed off NGM plates in S-media that contains 0.02% Triton. This allows for a more consistent number of worms while pipetting, as less worms stick to the plastic pipette tips. The volume is adjusted with S-media until there would be 60-70 worms per 20ul. Worms are scored as paralyzed if their body cannot make a bending “S” movement. Paralyzed worms can often still make small movements with their head or tail. Videos are captured using a Lumenera Infinity 3s camera fitted to a Nikon TE300 microscope at 2x magnification and recorded to ImageJ. In some experiments videos are captured using Phylumtech’s Wormtracker machine. The videos will be analyzed using ImageJ *C. elegans* motility analysis software. Level of activity will be denoted based on improvement in swimming induced paralysis (SWIP) in human TDP-43 transgenic *C. elegans* disease model. The automation data measuring paralysis will measure average body bends per second of a population. Improvement in SWIP from control in the population of worms will also be observed.

#### Drosophila Drug feeding assay

Melt cornmeal-molasses-yeast fly food was mixed with certain concentrations of compound at high temperature and cooled to RT. DMSO was used as the vehicle control. Parent flies were crossed on food supplemented with drugs and the offspring were raised on the same food. Adult flies were aged on the drug-containing food for 15 days before analyzing their eye morphology. For quantification of outer eye morphological defects, ten flies were quantified.

#### SODG93A Mouse Efficacy Study

Wildtype and SODG93A mutant mice were grown for 5 weeks, then given daily IP doses with vehicle, compound T18, or compound T20 for another 5 weeks. Weight and serum pNHF were tracked during the study.

## Abbreviations

(ALS): Amyotrophic lateral sclerosis
(rATP): Adenosine triphosphate
(BSA): Bovine serum albumin
(CFPSA): Cell-free protein synthesis and assembly
(rCTP): Cytidine triphosphate
(eDRAC): Energy-dependent drug resin affinity chromatography
(FTD): Fronto-temporal dementia
(rGTP): Guanosine triphosphate
(HIV): Human immunodeficiency virus
(IP): Intraperitoneal
(LC): Liquid chromatography
(MS): Mass spectrometry
(mL): Milliliter
(mM): Millimolar
(mg): Milligram
(PDF): Patient-derived fibroblasts
(PBS): Phosphate buffered saline
(pNFH): Phosphorylated neurofilament heavy chain
(PDI): Protein disulfide isomerase
(SAP): Streptavidin precipitation
(SG): Stress granule
(SAR): Structure-activity relationship
(SWIP): Swimming-induced paralysis
(MS-MS): Tandem mass spectrometry
(THIQ): Tetrahydroisoquinolone
(TDP-43): Transactive DNA-binding protein of 43 kDa
(UTP): Uridine triphosphate

## Acknowledgments

We thank the Target ALS postmortem tissue core for providing blinded ALS and healthy tissues. Funding for these studies was provided by Prosetta Biosciences.

