## Supplemental Figures for "Protein Assembly Modulation: A New Approach to ALS Therapeutics"

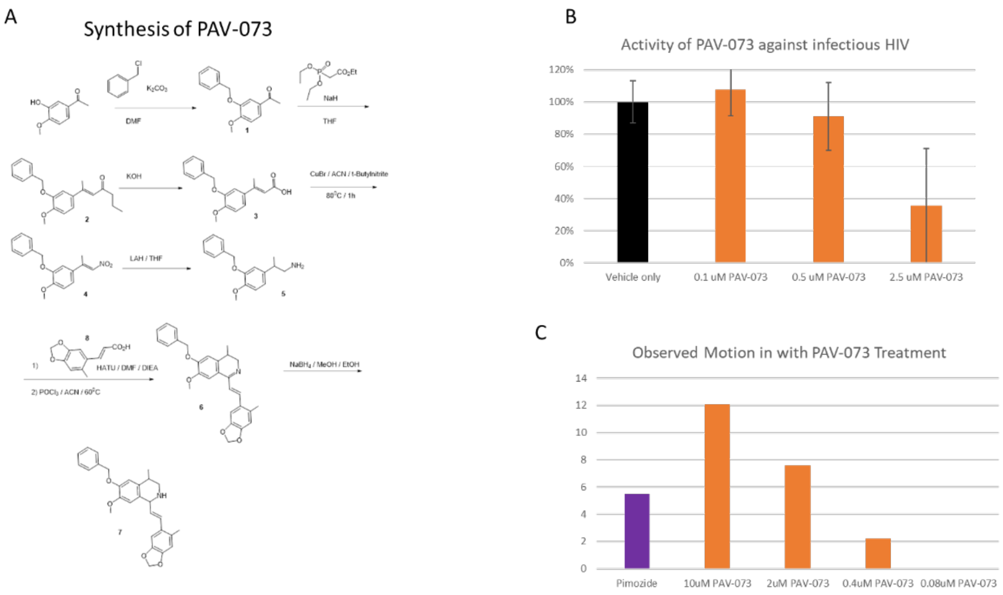


**Supplemental Figure 1. Synthesis and activity of PAV-073. Supplemental Figure 1A** shows the synthetic scheme for PAV-073. **Supplemental Figure 1B** shows activity of PAV-073 against infectious HIV. MT-2 cells were infected with NL4-3 Rluc HIV and treated with PAV-073 for four days. Averages and standard deviation of viral titer observed with triplicate repeated dose-titrations of PAV-073 are shown as a percentage of the titer observed in DMSO-treated cells. **Supplemental Figure 1C** shows activity of PAV-073 relative to pimozide (40uM) in ameliorating the condition of transgenic *C. elegans* expressing the human TDP-43 A315T mutation. Nematodes were age-matched and grown on standard nematode grown media plates until day 1 of adulthood at which point they were collected and placed in 96 well plates (50-70 animals per well) and treated with compound or control. Animal movement was then tracked for 30 minutes using WMicroTracker ONE.

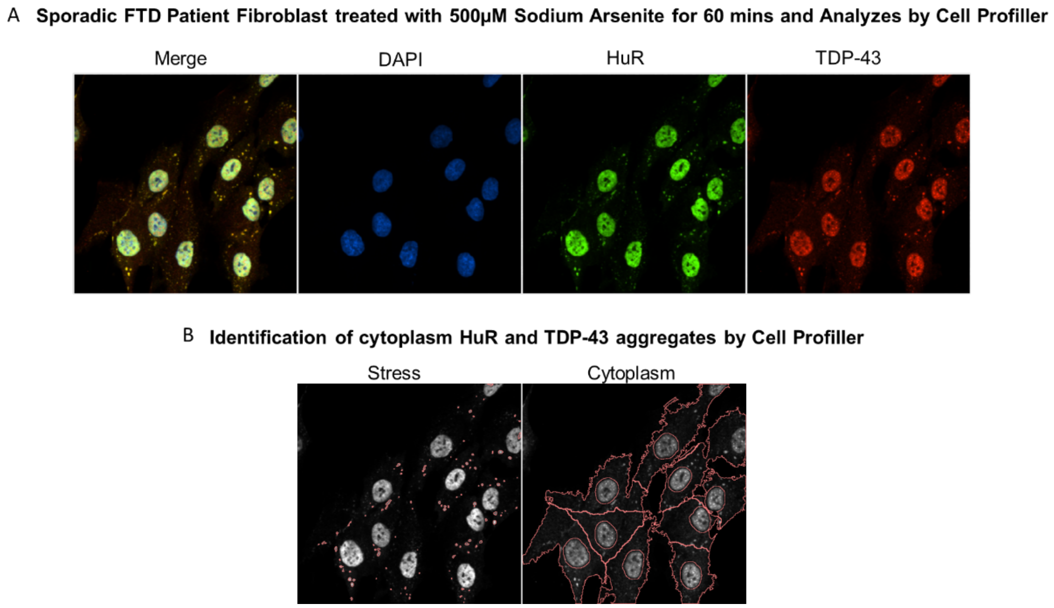


**Supplemental Figure 2. Stress granule induction and quantitation. Supplemental Figure 2A** shows immunostain for DAPI, HuR, and TDP-43 in PDFs treated with 500uM sodium arsenite for one hour. **Figure 2B** shows how cell profiler imaging was used to identify the number of TDP-43 positive HuR aggregates per cell.

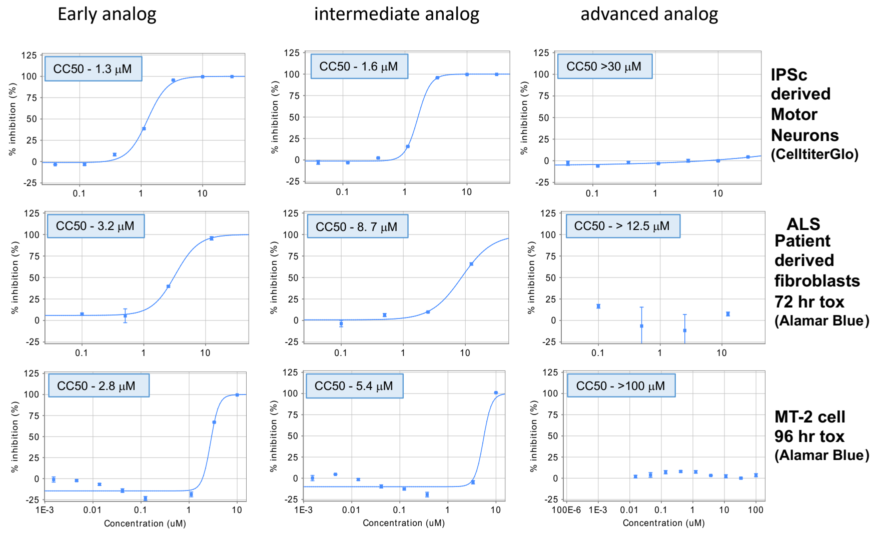


**Supplemental Figure 3.** Moderation of toxicity with THIQ lead series advancement. Compounds of comparable potency but differences in toxicity were identified from the lead series and assessed in transformed cells, ALS PDFs, and iPSC-derived motor neurons. The lead series progression to diminished toxicity is confirmed in all cell lines using both cell TiterGlo and Alamar Blue toxicity assays.

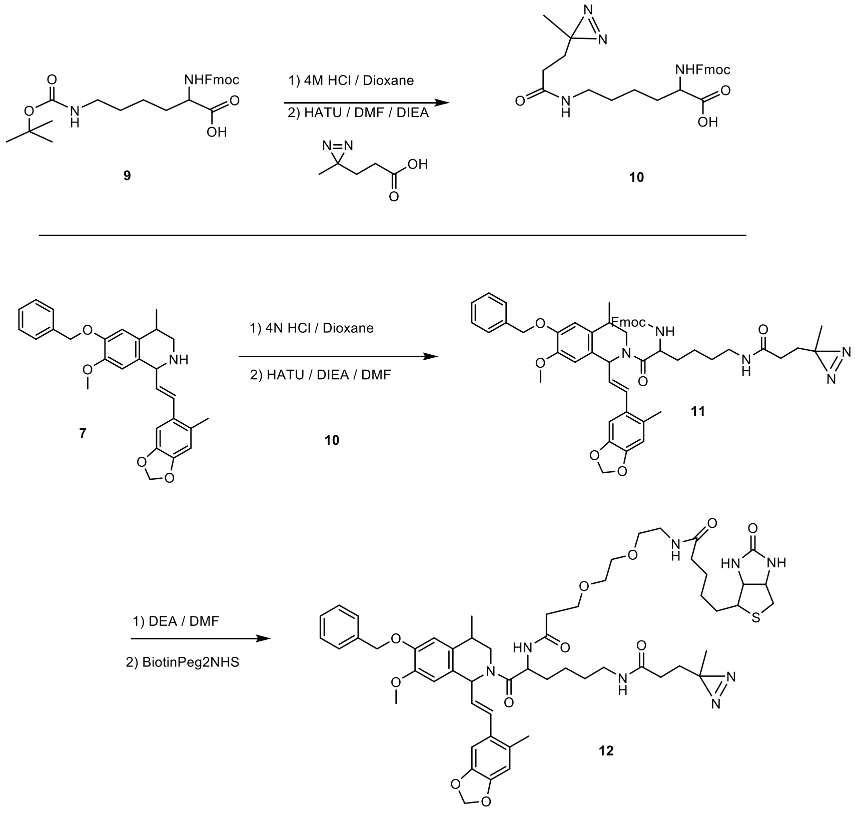


**Supplemental Figure 4. Synthetic scheme for PAV-073 photocrosslinker analog. Supplemental Figure 3** shows the synthetic scheme for the photocrosslinker analog of PAV-073 used to identify PDI as the direct drug binding protein (see **Figure 5**).
